# Cross-species transmission and PB2 mammalian adaptations of highly pathogenic avian influenza A/H5N1 viruses in Chile

**DOI:** 10.1101/2023.06.30.547205

**Authors:** Catalina Pardo-Roa, Martha I. Nelson, Naomi Ariyama, Carolina Aguayo, Leonardo I. Almonacid, Gabriela Munoz, Carlos Navarro, Claudia Avila, Mauricio Ulloa, Rodolfo Reyes, Eugenia Fuentes Luppichini, Christian Mathieu, Ricardo Vergara, Álvaro González, Carmen Gloria González, Hugo Araya, Jorge Fernández, Rodrigo Fasce, Magdalena Johow, Rafael A. Medina, Victor Neira

**Affiliations:** Department of Child and Adolescent Health, School of Nursing, Pontificia Universidad Católica de Chile, Santiago, Chile; Department of Pediatric Infectious Diseases and Immunology, School of Medicine, Pontificia Universidad Católica de Chile, Santiago, Chile; National Center for Biotechnology Information, National Library of Medicine, National Institutes of Health, Bethesda, Maryland 20892, USA; Departamento de Medicina Preventiva Animal, Facultad de Ciencias Veterinarias y Pecuarias, Universidad de Chile. 11735 Santa Rosa, La Pintana, Santiago, Chile; Servicio Agrícola y Ganadero, SAG, Chile; Molecular Bioinformatics Laboratory, Department of Molecular Genetics and Microbiology, Faculty of Biological Sciences, Pontificia Universidad Católica de Chile, Santiago, Chile; Institute for Biological and Medical Engineering, Schools of Engineering, Medicine and Biological Sciences, Pontificia Universidad Católica de Chile, Santiago, Chile; Servicio Nacional de Pesca y Acuicultura, SERNAPESCA, Chile; Veterinary Histology and Pathology, Institute of Animal Health and Food Safety, Veterinary School, University of Las Palmas de Gran Canaria, Las Palmas de Gran Canaria, Spain; Instituto de Salud Pública, ISP, Ministerio de Salud, Santiago, Chile; Department of Pathology and Laboratory Medicine, School of Medicine, Emory Vaccine Center, Emory University, Atlanta, USA; Department of Microbiology, Icahn School of Medicine at Mount Sinai, New York, USA

**Author notes:** Contributed equally (Catalina Pardo-Roa, Martha I. Nelson, Naomi Ariyama).

**Keywords:** H5N1, HPAIV, Panzootic, Marine Mammals, Chile, Genomic Surveillance, Interspecies transmission, Zoonotic

## Abstract

H5N1 highly pathogenic avian influenza viruses (HPAIV) emerged in wild birds in Chile in December 2022 and spilled over into poultry, marine mammals, and one human. Between December 9, 2022 – March 14, 2023, a coordinated government/academic response detected HPAIV by real-time RT-PCR in 8.5% (412/4735) of samples from 23 avian and 3 mammal orders. Whole-genome sequences obtained from 77 birds and 8 marine mammals revealed that all Chilean H5N1 viruses belong to lineage 2.3.4.4b and cluster monophyletically with viruses from Peru, indicating a single introduction from North America into Peru/Chile. Mammalian adaptations were identified in the PB2 segment: D701N in two sea lions, one human, and one shorebird, and Q591K in the human and one sea lion. Minor variant analysis revealed that D701N was present in 52.9 – 70.9% of sequence reads, indicating the presence of both genotypes within hosts. Further surveillance of spillover events is warranted to assess the emergence and potential onward transmission of mammalian adapted H5N1 HPAIV in South America.

## Introduction

Influenza A virus (IAV) is a segmented negative single-stranded RNA virus with a remarkable capacity to spillover and adapt to new host species, resulting in epidemics and pandemics ^1, 2^. Aquatic wild birds are considered the primary natural reservoir of IAV and harbor 16 HA subtypes and 9 NA subtypes. Endemic and migratory birds interact across established migratory flyways, which can serve as routes for long-distance IAV dispersal ^3^. Highly pathogenic avian influenza viruses (HPAIV) present a major threat to animal health, causing important losses to the poultry industry and affecting wild birds and occasionally mammal populations. Zoonotic cases of HPAIV of the H5N1 subtype have high fatality rates in humans, upwards of 50%, but have been concentrated in Asia thus far ^4^.

Since 2020, outbreaks caused by the influenza A H5N1 2.3.4.4b clade have been reported in Europe, Asia, Africa, and North America ^5^. In October 2022, HPAIV H5N1 emerged in Latin America ^6^ with cases identified in Peru, Venezuela, Chile, Mexico, and Ecuador ^6, 7^. To date, the reporting of HPAIV H5N1 in mammals, including humans, continues to be linked to high viral loads associated with the presence of positive wild birds ^8, 9^. Given the 2.3.4.4b virus’s potential for reassortment, efficient transmission in avian populations, and the ability to transmit across species, this lineage has recently been included in the World Health Organization’s list of candidate vaccine viruses for zoonotic influenza ^10, 11^.

The first case in Chile was confirmed in December 2022 ^12^, resulting in large outbreaks in seabirds. The virus rapidly disseminated across the country, covering approximately 2,300 kilometers over four months, from early December 2022 to March 2023. During this period, HPAIV affected a diverse array of species, including but not limited to seabirds, scavengers, penguins, raptors, marine mammals such as sea lions, backyard poultry, and intensive poultry flocks. In March 2023, a positive human case was confirmed ^13^. The patient developed severe viral pneumonia and was hospitalized requiring ventilation support. including cough, sore throat, and hoarseness and sought medical treatment at a local hospital on March 21 after symptoms progressed. The patient developed difficulty breathing (dyspnea) and was transferred to a regional hospital, where a bronchoalveolar sample was collected on March 27. The sample tested positive for influenza A virus by real-time RT-PCR by the Chilean Institute of Public Health (ISP) and was further characterized as H5N1 HPAIV by the WHO Collaborating Centre ^14^.

The high frequency of HPAIV spillover into mammals – and now a human – raises concerns that the virus could be evolving towards more efficient mammalian transmission and ultimately increase the potential of a global pandemic. To track the evolving threat of HPAIV H5N1 in Chile, we conducted comprehensive genomic surveillance of HPAIV H5N1 along Chile’s long Pacific coastline from December 2022 to March 2023. We employed real-time RT-PCR testing for initial detection and H5N1 confirmation, followed by whole genome sequencing of positive cases. Notably, our study represents a large national surveillance effort to sample suspected infected avian and mammalian species, including humans, and encompassing both wildlife and livestock sectors, to assess the spread of infection, impact on populations, and zoonotic risk. These findings contribute to improve our understanding of the evolutionary dynamics of HPAIV H5N1 and inform targeted strategies for disease management and control.

## Methods

### Sample collection

This study includes active and passive surveillance performed by the Chilean Agricultural and Livestock Service (SAG) and the National Fisheries Services (SERNAPESCA). Sampling was conducted between December 1, 2022, and March 14^th^, 2023, with the objective of tracing the HPAIV H5N1 outbreak across the country. Samples were collected from the Chilean territory, spanning approximately between parallels 18 and 54 south latitudes, according to the recommendation from the United States Department of Agriculture National Veterinary Services Laboratories (USDA-NVSL) protocol NVSL-WI-0023. Briefly, oropharyngeal (ORP), tracheal (TRS), and cloacal (CLO) swabs, feces, or tissues were collected from gallinaceous poultry (ORP or TRS), domestic waterfowl and other wild birds (ORP or CLO). TRS, rectal swabs (RCS), and nasal swabs (NAS) or spiracle swabs from marine mammals were collected on cryotubes containing viral transport media (NVSL media #10088 10%, P/S 1% Gentamicin 0,1%, and Fungizone) and were then submitted for diagnostic reverse transcription PCR (RT-PCR) to the reference laboratory. Pools of up to 10 combined cloacal/tracheal (CCT) or CLO swabs were processed from gallinaceous poultry and domestic ducks, which were grouped by the same species, the same premises, and the same sampling route, in accordance with NVSL guidelines.

A total of 9,980 samples, obtained from 4,737 submission cases, were derived for viral diagnostic. Of them, 61.7% of samples were from poultry (n=6,156), 37.1% from wildlife birds (n=3,708), and 1.2% from non-human mammals (n=116). Avian samples were collected from 26 different orders, including highly represented orders such as, *Galliformes* (47.89%)*, Charadriiformes* (15.13%), *Anseriformes* (7.32%) to lesser represented order, such as *Coraciiformes,* and *Tinamiformes* (0.02% each). Also, non-human mammal samples from orders *Artiodactyla* (Burmeister’s porpoise n=1); *Cetartiodactyla* (Megaptera novaeangliae n=1) and *Carnivora* (South America sea lion n=67, marine otter n=4 and canines n=2;) were tested (**Supplementary information Table 1**).

### Molecular Diagnosis of Influenza A Virus

The diagnosis was performed at the SAG Livestock Virology Laboratory and the Biotechnology Laboratory (at SAG, Lo Aguirre). First, RNA was extracted from samples by the MagMAX™ Core Viral/Pathogen kit (Thermofisher, AM1830) using the KingFisher ThermoScientific™ KingFisher™ Flex Purification Systems. Then, real-time RT-PCR for AIV diagnosis was done using the VetMAX-Gold AIV Detection Kit (cutoff Ct value = 38 / internal control 25-30 Ct, Applied Biosystems™ Cat No. 4485261) and the Detection Kit Genome amplification (Thermo Fisher Scientific, internal control 25-30 Ct). Positive samples were tested to determine the Influenza A subtype H5 lineage (specific 2.3.4.4 clade) with specific real-time RT-PCR following the standard operating procedures of NVSL-USDA ^15–17^

### Whole genome Sequencing

Sequencing of positive H5 HPAIV samples was performed at the Laboratory of Molecular Virology, Pontificia Universidad Católica de Chile. To guarantee the amplification of all viral segments, viral RNA was re-extracted from the original sample with TRIzol^TM^ lysis (Invitrogen ^TM^ 15596018) followed by extraction using E.Z.N.A Viral RNA Kit (R6874, Omega Biotek). Then, the viral genome was amplified using a multi-segment one-step RT-PCR genome amplification with primers, Opti1-F1 5’:GTTACGCGCCAGCAAAAGCAGG-3’, Opti1-F25’:GTTACGCGCCAGCGAAAGCAGG-3’ and Opti1-R1:5’:GTTACGCGCCAGTAGAAA-CAAGG-3’ (13), followed by the amplicon purification with SPRISelect Beads (Beckman counter B2338) using a ratio of 0,45x. Then, next-generation sequencing was performed by Oxford Nanopore Technology (ONT) using the Native Barcoding Kit (SQK-NBD114.96) for library preparation. First End-prep was performed using the NEBNext Ultra II End Repair / dA-tailing Module (NEB, E7546), the native barcodes were ligated with the Native Barcoding Expansion 96 (EXP-NBD196) and NEB Blunt/TA Ligase Master Mix (NEB, M0367) selecting a unique barcode per sample. The barcoded library was pooled together and purified using SPRISelect beads. An AMII adapter was ligated to the library using the NEBNext Quick Ligation Module (NEB, E6056), purified using SPRISelect beads, and quantified with a Qubit Fluorometer (Invitrogen ^TM^). The library was loaded on the sequencer using the ligation sequencing kit (SQK-LSK109) according to ONT instructions for the R.9 flow cells. Sequencing was carried out for 72 hours.

Nanopore Reads (FASTQ) were filtered with NanoFilt requiring an average quality of at least 7 and a length not exceeding 2600 nucleotides ^18^. Genomes were assembled by reference using the Nanopore ARTIC pipeline (v1.2.3) and modified appropriately ^12^. Each assembled segment was then used to search for the most similar influenza segment within the Influenza Virus Resource database (IRD) of NCBI, to then use it as a template for an assembly by reference. The reference was chosen by BLAST-searching preliminary assembled contigs constructed with filtered reads which were de novo assembled with Canu ^19^. Reads were mapped to the selected reference with minimap2 and used to build a consensus sequence for each segment, which was carried out with the tools medaka, longshot, Samtools y Bcftools ^20–22^. Assembly of IAV genomic segments was performed using a custom pipeline in multiple stages. First, an initial assembly was done using the inchworm component of Trinity, and viral contigs bearing internal deletions will be identified by BLAST mapping against non-redundant IRD reference sequences ^23, 24^. In the second stage the inchworm assembly was repeated but now removing breakpoint-spanning kmers from the assembly graph. The resulting IAV contigs were then oriented and trimmed to remove low coverage ends and any extraneous sequences beyond the conserved IAV termini. In the final stage, the CAP3 assembler was used to improve contiguity by merging contigs originating from the same segment type if their ends overlapped by at least 25 nt ^25^. Once consensus genomes were obtained, they were checked for quality and annotated by the NCBI Influenza Virus Sequence Annotation Tool (https://www.ncbi.nlm.nih.gov/genomes/FLU/annotation/) ^26^. H5 clade classification was carried out with the online tool Subspecies Classification at the BV-BRC portal ^27, 28^. One hundred twelve samples were selected and processed and eighty-five of them were successfully amplified and sequenced. Sixty-nine complete genomes and ten partial genomes were obtained (with at least 6 segments, **Supplementary information Table 2**). All sequences have been deposited in GenBank (Submission ID: 1136917325075).

### HPAIV H5N1 human case

On March 13, 2023, a 53-year-old male from the Region of Antofagasta in northern Chile began experiencing symptoms, including cough, sore throat, hoarseness, and sought medical treatment at a local hospital on March 21 after symptoms progressed. The individual had no declared comorbidities, recent travel history or visit to beaches. Subsequently, on 22 March 2023, the patient developed difficulty breathing (dyspnea) and was transferred to a Regional Hospital in Antofagasta, which serves as a severe acute respiratory infection (SARI) Sentinel Site. A nasopharyngeal swab sample was collected as part of routine SARI surveillance and tested negative for SARS-CoV-2 using RT-PCR. On 23 March, the patient was transferred to the intensive care unit, where antiviral treatment with oseltamivir and antibiotics was initiated. A bronchoalveolar sample collected on 27 March tested positive by RT-PCR for a non-subtyped influenza A virus. Subsequent subtyping by real-time RT-PCR confirmed avian influenza A(H5) on 29 March and the Ministry of Health of Chile (MINSAL) reported the first case of infection with avian influenza A (H5) virus in Chile to the World Health Organization ^29^. The patient’s samples were forwarded to a WHO Collaborating Centre for further characterization. On April 5^th^, genomic sequencing conducted by the National Institute of Public Health in Chile confirmed it as HPAIV H5N1 clade 2.3.4.4b. Sequences were deposited in GISAID Accession numbers EPI2510180-EPI2510187. As of June 16, 2023, the patient remains in respiratory isolation and requires mechanical ventilation due to pneumonia.

All three close contacts of the patient were identified, did not present clinical signs and tested negative for influenza. They completed the monitoring period successfully. Among healthcare workers, a total of nine contacts were identified, all of whom tested negative for influenza and concluded their monitoring period on April 4^th^. However, on April 5^th^, one healthcare worker developed respiratory symptoms. A nasopharyngeal swab test yielded a negative result for influenza and any other respiratory virus. The monitoring period for the contact was extended for an additional seven days, ending on April 11^th^, 2023. Importantly, HPAI had previously been identified in wild aquatic birds (pelicans and penguins) as well as sea mammals (sea lions) in the Antofagasta Region during the period of December 2022 to February 2023. Initial findings from the epidemiological investigation suggest that the most probable mode of transmission for this human case was through environmental exposure, considering the significant presence of deceased sea mammals and wild birds near the patient’s residence, which is within 150 meters of the beach.

### Phylogenetic analysis

To place the newly obtained Chilean HPAIV H5N1 sequences in a global context, we downloaded a global background data set of all available AIV sequences for all segments from GenBank and GISAID databases that were submitted January 1, 2021 – April 27, 2023, including avian and mammalian hosts, and aligned them with the Chilean sequences generated in this study, including the human case, using MAFFT ^30^. Larger datasets (ranging from 5,253 - 5,616 sequences) were used for internal gene segments, which included all AIV subtypes. Smaller datasets were used for HA and NA segments since these were limited to one subtype: H5 (n = 1,058 sequences) and N1 (2,534 sequences). Phylogenetic trees were inferred for each segment separately using maximum likelihood (ML) methods available in IQ-Tree2 ^31^ with a GTR model of nucleotide substitution, incorporating gamma-distributed rate variation among sites. To assess the reliability of each node in the phylogenetic trees, a bootstrap resampling process was performed with 1000 replicates. Due to the size of the dataset, we used the high-performance computational capabilities of the Biowulf Linux cluster at the National Institutes of Health (http://biowulf.nih.gov).

After the initial ML trees showed that all Chilean viruses clustered with Peru viruses in a single clade on all eight gene segment trees, an additional phylogenetic analysis was performed on just this clade for all eight segments using a Bayesian phylogeographic approach ^32^. We used the Markov chain Monte Carlo (MCMC) method available in the BEAST package, v1.10.5 ^33^, again using the NIH Biowulf Linux cluster. An exponential demographic model was used in this outbreak setting, with a general-time reversible (GTR) model of nucleotide substitution with gamma-distributed rate variation among sites. Each tip was assigned a location state and a phylogeographic discrete trait analysis was performed. The MCMC chain was run separately four times for each dataset using the BEAGLE 3 ^34^ library to improve computational performance, until all parameters reached convergence, as assessed visually using Tracer v.1.7.2 ^35^. At least 10% of the chain was removed as burn-in, and runs for the same dataset were combined using LogCombiner v1.10.4. A MCC tree was summarized using TreeAnnotator v.1.10.4 and visualized in FigTree v1.4.4. The analysis was repeated for six of the eight genome segments (PB2, PA, HA, NP, NA, and MP) for which the majority of viruses in the Peru/Chile clade had sequences available and showed no evidence of reassortment (n = 79). When mammalian adapted PB2 mutations were identified in Chilean samples, an additional phylogenetic analysis was performed to study the evolutionary relationships of viruses with these PB2 mutations. The analysis used similar ML methods as described above and an updated dataset of all H5N1 PB2 segments submitted to GISAID January 1, 2021 – June 8, 2023, including avian and mammalian hosts (n = 4,120 H5N1 PB2 sequences). To visualize the spatial dispersal of H5N1 viruses in North America, we performed a continuous phylogeographic analysis, again implemented in BEAST 1.10.5, assigning a geographical coordinate (latitude and longitude) to each tip and using a Cauchy distribution to model among-branch heterogeneity in diffusion velocity ^36^. MCMC chains were run for at least 200 million iterations, to reach adequate ESS values as estimated by Tracer 1.7, discarding 10% of sampled trees as burn-in. SPREAD 4 was used to visualize continuous dispersal events from the MCC tree ^37^. All XML and tree files are publicly available in the GitHub repository (https://github.com/mostmarmot/ChileHPAIV).

To identify specific genetic variations and mutations that may have implications for the pathogenicity, transmission, and antigenic characteristics of the H5N1 viruses under investigation, we conducted mutation analysis using the CDC H5N1 genetic changes inventory for SNP analysis and incorporated various other previously published mutations of concern ^27, 38^.

### Mutational and minor single variants analysis

To assess the presence of mutations and minor variants of the sequenced genomes, the raw data (FASTQ) was filtered with NanoFilt ^18^, keeping reads with an average quality of ≥ 7 (Phred score) and a length between 400 and 2,600 bp. Filtered reads were then mapped to the influenza A (H5N1) reference genome with Minimap2 ^22^. The reference used was the first case of avian H5N1 Influenza virus, recently reported in Chile, the strain A/black skimmer/Chile/C61962/2022, GenBank OQ352545-OQ352552. Unaligned reads and supplementary alignments were removed from BAM files with Samtools ^21^. Single nucleotide variant (SNV) and indel calling was carried out with Clair3, configured for haploid genomes, enabling the “haploid_sensitive” option and setting 0.5% and 5% as minimum SNV and indel frequency threshold, respectively ^39^. Called variants were then annotated with SnpEff, including options “-no-downstream” and “-no-upstream” ^40^. Finally, variants were plotted with Circos ^41^. This analysis did not include the human case.

## Results

### HPAIV H5N1 outbreak in Chile

Out of 4,737 submitted cases, 412 were positive by real-time RT-PCR (8.45%) and were later confirmed and subtyped as H5N1. The 412 H5N1 cases were distributed across a 3,000+ kilometer stretch of Chile’s Pacific coastline extending from the Arica Region in northern Chile (Lat: −18.419 long: −-70.3212) to the Los Lagos Region in southern Chile (Lat: −41.866 Long: −73.8290) (Figure 1A). Positive cases belonged to 33 different animal species, including birds and mammals, wildlife and domestic, and backyard and commercial poultry. Thirty-one cases belonged to poultry and 30 of them corresponded to backyard farms and one case of a broiler commercial breeding flock. Backyard-positive animals included chickens, turkeys, domestic ducks, and geese. Three hundred and sixty-three cases belonged to avian wildlife. Of these, 113 (27,43%) corresponded to Pelicans (*Pelecanus thagus*) samples, 70 (16,99%) were Peruvian booby (*Sula variegata*), 50 (12,14%) were Guanay shag (*Leucocarbo bougainvilliorum*), 31 (7,42%) were kelp gull (*Larus dominicanus*) and 49 (11,89 %) were from other gulls. Additionally, positive cases were detected in vultures (n=19), swans (n=3), and Humboldt penguins (n=4). Eighteen positive cases belonged to marine mammals: 16 were from South American sea lions *(Otaria flavescens*) and two were from endangered marine otters (*Lontra felina)*; a complete list of positive cases is in **Supplementary Information Table 1 and 3)**. Positivity rates were highest in *Pelecaniformes* (48,3%), *Podicipediformes* (33,3%), *Cathartiformes* (29,6%), *Suliformes* (28,2%) *and South America sea lions* (23.8%) (**Supplementary Table 1**). All sampled marine mammals presented unspecific signs of disease such as weakness, anorexia, and weight loss, and other specific ones such as acute dyspnea, tachypnea, profuse nasal secretion, sialorrhea, and abdominal breathing (respiratory syndrome), as well as neurological signs such as tremors, ataxia, paralysis of limbs, disorientation, and nystagmus (neurological syndrome). An estimated 2400% increase (24 times) in the historical marine mammals strandings and death have been observed since the first detection of HPIAV H5N1 in Chile (SERNAPESCA national records).

**Figure 1.**
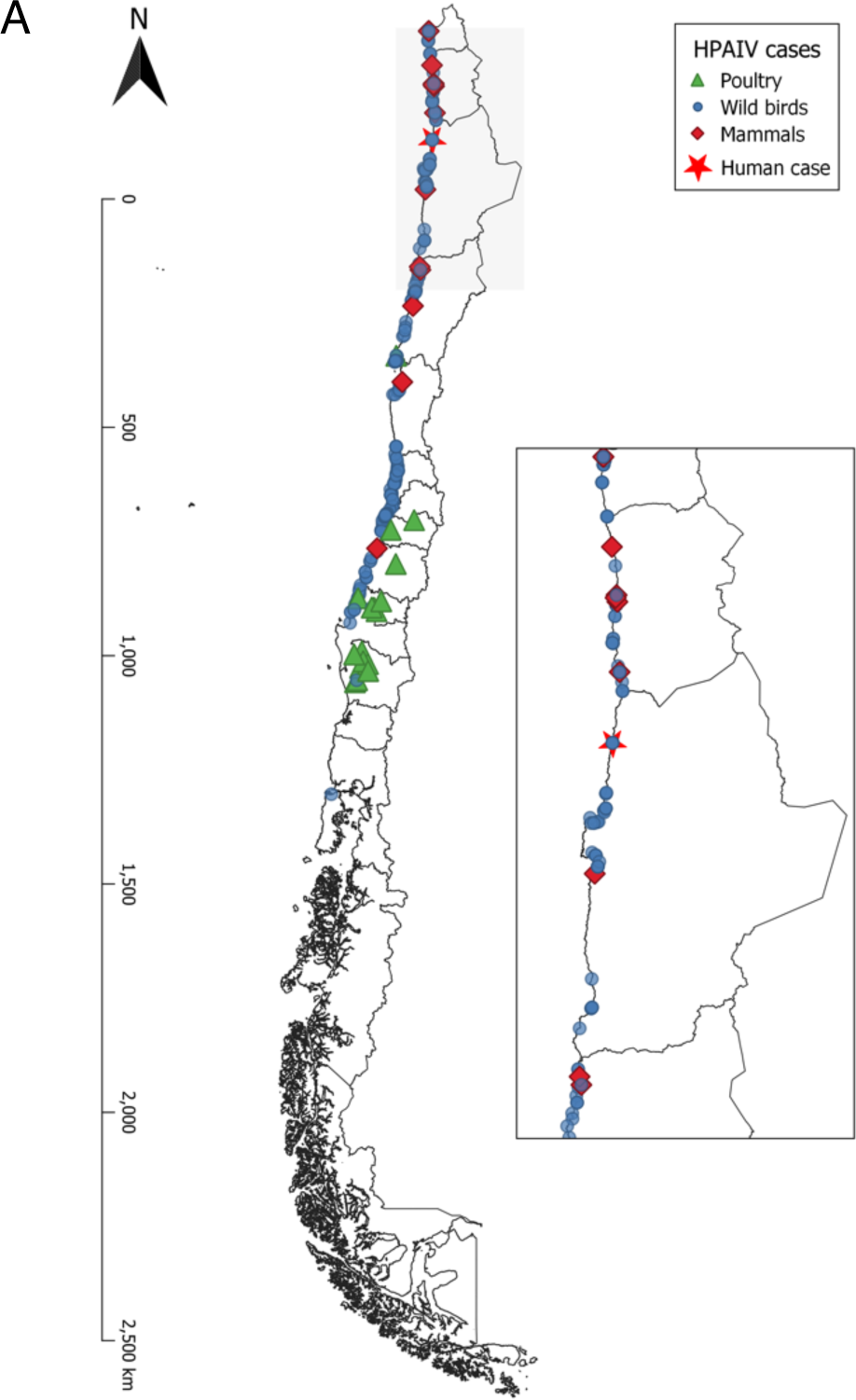
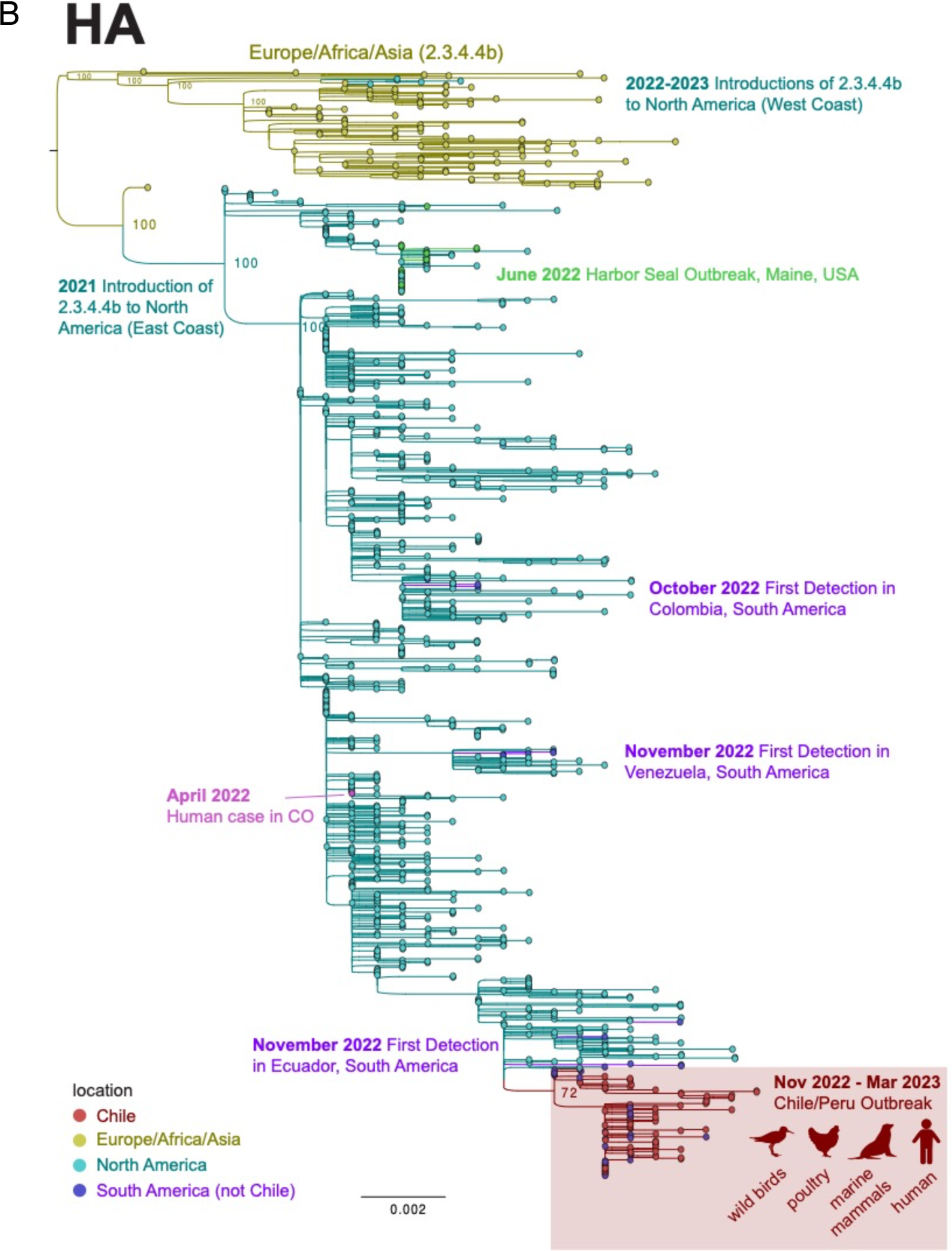
Geographical distribution and phylogenetic relationships of HPAIV/H5N1 2.3.4.4b viruses in Chile and globally. **A.** Distribution of positive samples collected between December 1, 2022, and March 14^th^, 2023. Inset depicts an amplified image of northern Chile where positive wild birds and marine mammal cases were identified near the HPAIV H5N1 human case. **B.** Maximum likelihood tree inferred for 1058 HA sequences from H5N1 viruses belonging to the 2.3.4.4b lineage that were collected globally from January 1, 2021 – March 24, 2023. Tree is midpoint rooted and bootstrap values are provided for key nodes. Tips and branches are shaded by geographical location. The clade of viruses from Chile and Peru collected during the November 2022-March 2023 outbreak is highlighted in a red box, with host species listed.

### Single Introduction of HPAIV H5N1 clade 2.3.4.4b into Chile

Phylogenetic analysis of all eight genome segments of H5N1 viruses collected from Chile have 4:4 reassortant genomes similar to the H5N1 viruses described recently in Peru ^42^. The reassortant viruses have retained four segments (PA, HA, NA, and MP) from the original Eurasian AIV lineage and acquired four new segments (PB2, PB1, NP, NS) from the Americas AIV lineage. The viruses from Chile and Peru cluster together into a single monophyletic cluster on all eight trees inferred for each genome segment (Figure 1B), representing a single viral introduction from North America into Peru/Chile. The trees also identified additional independent introductions of H5N1 viruses from North America into other South American countries, including Colombia, Ecuador, and Venezuela (see http://bit.ly/3Jye2FN for an animated visualization inferred from the MCC tree). No transmission between these other South American countries was evident; rather, each country acquired its H5N1 population independently from North America (**Figure 1**). Only one full-length NA sequence from an H5N1 virus was available from Argentina, and this virus is positioned in the Peru/Chile cluster, suggesting continental spread of H5N1 to Argentina by way of Peru/Chile, based on this limited data.

### Spatial dispersal of HPAIV H5N1 in Chile

To study the spatial dispersal of H5N1 viruses along Chile’s thousands of kilometers of coastline in finer detail, a MCC tree of the Peru/Chile cluster was inferred from a concatenated alignment of six of eight genome segments, using a phylogeographic approach (**Figure 2A**). Chile’s H5N1 population formed two discrete clades that emerged in late 2022: a ‘Central Chile’ clade that includes poultry viruses from central Chile’s outbreaks in February-March 2023 and a larger “Major” clade that spans a longer period (December 2, 2022 – March 6, 2023), more diverse avian and mammalian host species, and a wider geographical range spanning from central Peru to central Chile. In the central Chile clade, poultry viruses were interspersed with wild bird viruses (e.g., mallard duck, black skimmer, great egret), indicating that poultry outbreaks affecting different regions in central Chile involved independent H5N1 spillovers from wild birds to poultry (**Figure 2B**). In the major clade, viruses from South American sea lions in Chile and Peru were also not monophyletic, consistent with multiple independent spillover events from wild birds to sea lions. In the major clade, viruses also were not monophyletic by region, consistent with regular viral gene flow between Chile’s regions, and between Peru and Chile.

**Figure 2.**
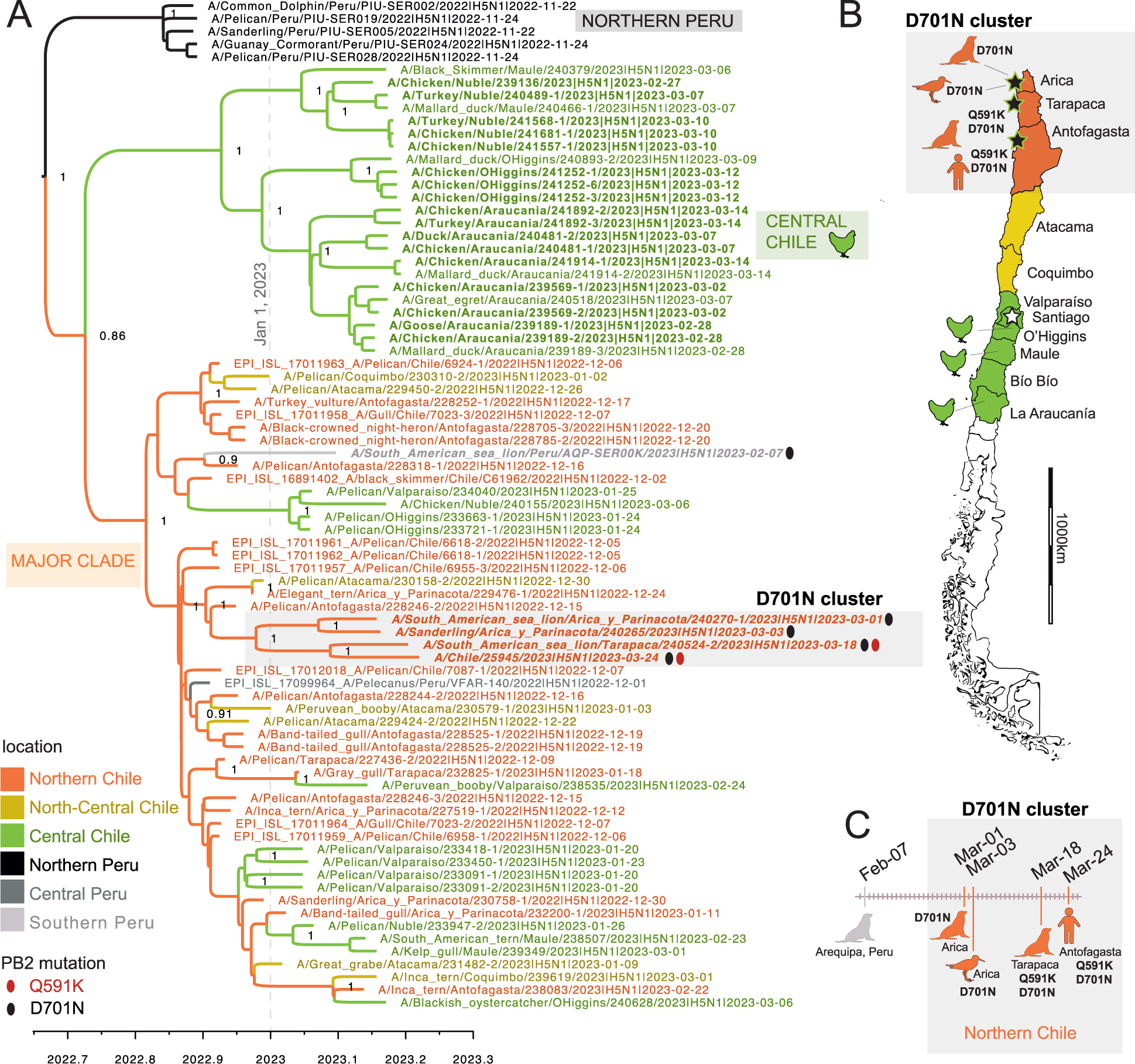
Phylogenetic relationships of the Chile/Peru HPAIV cluster. (A) Time-scaled MCC tree inferred for the partially concatenated genomes (∼10 kb) of 79 H5N1/2.3.4.4b viruses from the Chile/Peru HPAIV cluster (see Figure 1), using a Bayesian phylogeographic approach. Branches are shaded by inferred location (regions in Peru and Chile, see Figure 2B). Posterior probabilities are provided for key nodes. Ovals indicate viruses with the Q591K (red) or D701N (black) amino acid substitution in the PB2. Three clades are labeled: Northern Peru, Central Chile, and Major Clade. (B) Map of Chile including locations of viruses from the D701N cluster (Figure 2A) and from poultry. (C) Timeline of identifications of viruses with Q591K and D701N mutations in Peru and Chile. (Note that one Peru sea lion virus with mutations that appears in Figure 3 is omitted from this figure because a complete genome sequence was not available.)

### PB2 mutations associated with mammalian adaptation

Two mutations in PB2 (Q591K and D701N) that have been previously associated with AIV adaptation to mammalian hosts were identified in Chile (**Figures 2 and 3**). D701N was observed in four Chilean viruses: the human case, two sea lions, and one shorebird (sanderling). Two viruses with D701N also had a Q591K substitution (the human and one sea lion). The four Chilean viruses with D701N were collected in different host species, but otherwise had similar features: (a) clustering together on the PB2 tree (**Figure 3A**) and concatenated genes tree (**Figure 2A**), (b) sharing additional common mutations in other genes (Supplement), (c) were from northern Chile (**Figure 2B**), and (d) were collected in March 2023 (**Figure 2C**). Two viruses from sea lions in southern Peru also have the D701N mutation. One Peru sea lion virus with D701N had sufficient genome coverage to be included in the concatenated genes tree (**Figure 2A**) but was collected a month earlier than the Chilean viruses (on February 7, 2023) and is positioned in a different section of the tree and represents an independent D701N mutation event. The second Peru sea lion virus with D701N was collected the same month as the Chilean sea lion viruses (March 2023) and clusters together with them on the PB2 tree (**Figure 3A**, posterior probability = 0.99). Within this Chile/Peru cluster, two viruses only have the D701N mutation (the sanderling and one sea lion) and three viruses have both the D701N and Q591K mutations (human and sea lions from Chile and Peru). The three viruses with Q591K cluster together in a well-supported sub-clade (posterior probability = 1).

**Figure 3.**
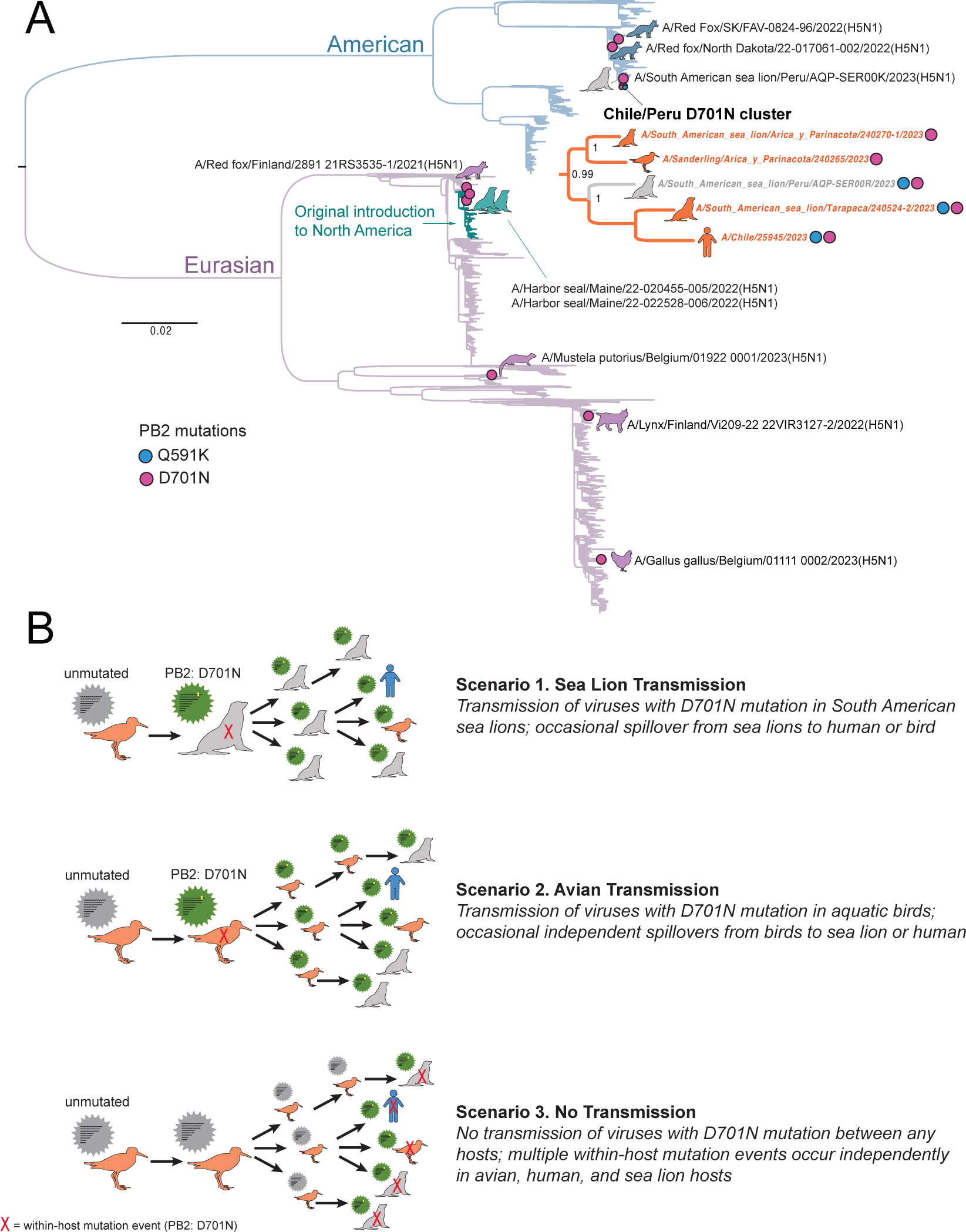
Evolutionary relationships of H5N1 viruses with mammalian adapted PB2 mutations. (A) Maximum likelihood tree inferred for 4120 PB2 sequences from H5N1 viruses collected globally in avian and mammalian hosts during January 1, 2021 – March 24, 2023. Branches are shaded by AIV lineage (American or Eurasian), with the recent introduction of Eurasian H5N1 into North America shaded teal. Pink circles with accompanying animal cartoons indicate viruses with D701N mutations; blue circles indicate Q591K. A detailed sub-tree is provided for the Chile/Peru D701N cluster. (B) Three scenarios for D701N transmission or *de novo* emergence.

For additional background context, the frequency of the Q591K and D701N mutations in H5N1 viruses circulating globally in other avian and mammalian populations was assessed among a larger dataset (n = 4,008 PB2 sequences) of H5N1 viruses collected globally since January 1, 2021. The D701N mutation was periodically observed in one-off events (i.e., no clusters) in four species of mammals (red fox (n = 3), harbor seal (n = 2), one lynx, one European polecat) and one bird (chicken), for a total global frequency of ∼0.2% (8/4008), not including Peru and Chile. The Q591K mutation was not observed in any H5N1 viruses outside of Peru and Chile (0/4008). In general, the Q591K and D701N mutations are exceedingly rare outside of Peru and Chile during the current global H5N1 epizootic. Moreover, the D701N occurrences appear to arise from *de novo* mutation events during replication in each individual host, with no observed onward transmission of mutated viruses between hosts. A central question for this study is whether the Q591K and D701N mutations observed in the Chile/Peru cluster are similarly *de novo* mutations that emerged independently in each host, or whether mutated viruses are transmitted between hosts, either in birds or marine mammals. At this time, multiple transmission scenarios involving wild birds, marine mammals, and humans could potentially explain the clusters of viruses with Q591K and D701N mutations in Chile/Peru (**Figure 3B**, see Discussion). Additional sequence data is needed from avian and mammalian hosts in Chile to fill in gaps in the tree and distinguish which scenario best fits the data.

### Minor variant analysis

In addition to the PB2 mutations Q591K and D701N, additional amino acid changes were observed in the larger Peru/Chile clade that were not associated with known phenotypic changes, including PB2: I616V; PB1: E264D, G399D, H5: L122Q (H3 numbering), NP: F230L, N1: L269M, S339P, MP: N85S, N87T and NS: C116S. To study within-host diversity, a minor single nucleotide (SNV) variant analysis was performed on the raw data of all the animal sequences, and several single nucleotide variants and inlets were found due to the compactness of the segments, most of the variations are related to gene sites that encode for a viral protein. A mean of 32,96 variations was found per genome, mainly associated with SNVs located on the polymerase genes (**Supplementary information Table 4**). Variations were classified according to the allele effect as low (synonymous_variant), moderate (missense_variant), high (frameshift_variant or stop_gained), or modifier (intron_variant or intergenic_region) **(Supplementary Table 5)**. The three samples located on the Chilean mammalian cluster (A/Sanderling/Arica y Parinacota/240265/2023(H5N1); A/South American sea lion/Arica y Parinacota/240270-1/2023(H5N1) and A/South American sea lion/Tarapaca/240524-2/2023(H5N1)) carry the D701N mutation at allele frequencies: 70,88%, 65,19% and 52,94; respectively. Nineteen additional SNVs were found on the PB2 D701N cluster located across the genome (**Figure 4**), eight of them were related to the cluster: PB2:D701N, HA: I194I, NP: I119T and C223C; M: H222H (with an intron variant) and NS: L69L and I226T (**Table 1**), suggestive of common genomic features of these viruses. Additional, viruses from marine mammal species would be needed to further the potential of onwards transmission of the D701N viruses.

**Figure 4.**
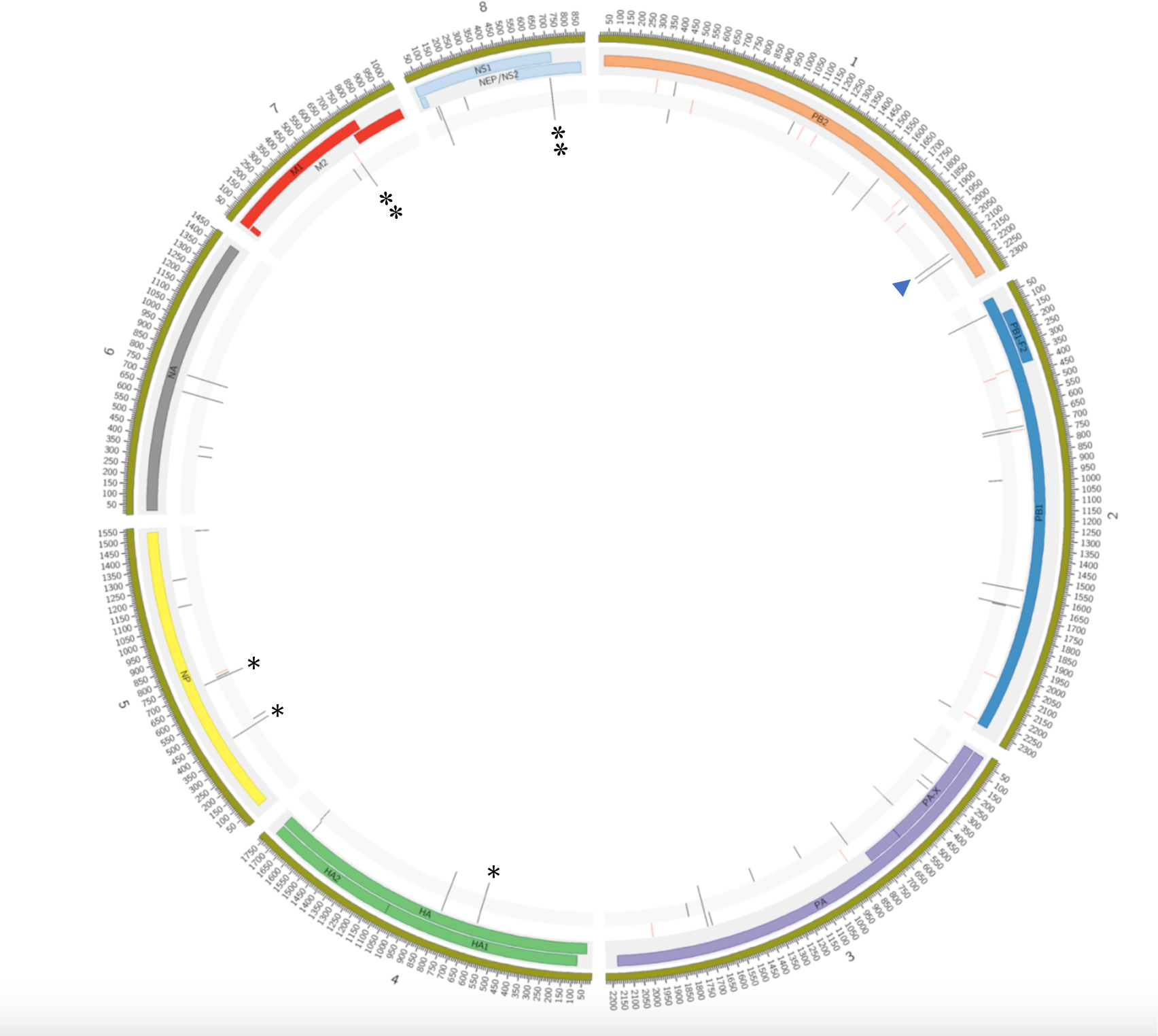
PB2 D701N clade shares other mutations throughout the viral genome. Nineteen SNVs were found in the 3 samples located on the PB2 D701N clade, seven of them related with mammal sequences (asterisks). Concentric circles represent, from the inside to the outside, the variants (red and black lines) found for samples A/Sanderling/Arica y Parinacota/240265/2023(H5N1), A/South American sea lion/Arica y Parinacota/240270-1/2023(H5N1) and A/South American sea lion/Arica y Parinacota/240270-1/2023(H5N1). Next, the viral proteins and their positions is annotated along the genome. The variations with a frequency <50% are marked in red and those with a frequency ≥ 50% are marked in black. The PB2-D701N mutation (arrowhead), present in all 3 samples, is located at genomic position 2.128 of segment 1. Red colored lines indicate the presence of the SNVs at frequencies of < 50% and black lines represent SNVs with >50% frequencies.

**Table 1:** Summary of minor variant mutations analysis of PB2 D701 Cluster sequences

| Chromosome / Segment | Genome position | Reference allele | Alternative allele | Alternative allele frequency / Position (Annotate allele) |  |  | Allele effect | Allele impact | CDS variation | CDS length | Protein variation | Comments |
| --- | --- | --- | --- | --- | --- | --- | --- | --- | --- | --- | --- | --- |
|  |  |  |  | 240524-2/2023 | 240265/2023 | 240270-1/2023 |  |  |  |  |  |  |
| OQ352545.1 / Seg 1 (PB2) | 1590 | T | A | 78,32 / 535 (A) | 76,92 / 65 (A) | 75,61 / 123 (A) | synonymous_variant | LOW | 1563T>A | 2280 | T521T | Present on 78 sequences |
|  | 2128 | G | A | 52,94 / 527 (A) | 70,88 / 1092 (A) | 65,19 / 135 (A) | missense_variant | MODERATE | 2101G>A | 2280 | D701N | Only on D701N cluster sequences |
|  | 2160 | T | C | 65,08 / 524 (C) | 87,35 / 1289 (C) | 90,37 / 135 (C) | synonymous_variant | LOW | 2133T>C | 2280 | N711N | Present on 39 sequences |
| OQ352546.1 / Seg 2 (PB1) | 96 | A | G | 100 / 3 (G) | 73,53 / 170 (G) | 86,15 / 65 (G) | synonymous_variant | LOW | 69A>G | 2274 | P23P | Present on 78 sequences |
|  | 705 | A | G | 100 / 3 (G) | 78,46 / 65 (G) | 85,94 / 65 /G) | synonymous_variant | LOW | 678A>G | 2274 | T226T | Present on 78 sequences |
|  | 723 | G | A | 33,33 / 3 (A) | 78,12 / 64 (A) | 84,38 / 64 (A) | synonymous_variant | LOW | 696G>A | 2274 | E232E | Present on 78 sequences |
|  | 1570 | T | G | 75 / 4 (G) | 76,27 / 118 (G) | 81,82 /154 (G) | missense_variant | MODERATE | 1543T>G | 2274 | S515A | Present on 40 sequences |
| OQ352547.1 / Seg 3 (PA) | 150 | A | G | 77,39 / 115 (G) | 77,39 / 115 (G) | 79,37 / 63 (G) | synonymous_variant | LOW | 126A>G | 2151 | L42L | Present on 33 sequences |
|  | 1692 | A | G | 73,39 / 124 (G) | 70,63 / 126 (G) | 73,13 / 67 (G) | synonymous_variant | LOW | 1668A>G | 2151 | Q556Q | Present on 37 sequences |
|  | 150 | A | G | 77,39 / 115 (G) | 77,39 / 115 (G) | 79,37 / 63 (G) | synonymous_variant | LOW | 126A>G | 759 | L42L | Present on 37 sequences |
| OQ352548.1 / Seg 4 (HA) | 610 | T | C | 89,97 / 3130 (C) | 88,94 / 461 (C) | 88,19 / 415 (C) | synonymous_variant | LOW | 582T>C | 1704 | I194I | Present on D701 cluster and 3 more samples sequences |
| OQ352549.1 / Seg 5 (NP) | 401 | T | C | 79,47 / 5255 (C) | 76,75 / 1045 (C) | 77,12 / 813 (C) | missense_variant | MODERATE | 356T>C | 1497 | I119T | Present on D701 cluster and 4 more samples sequences |
|  | 714 | C | T | 63,24 / 5283 (T) | 60,36 / 1047 (T) | 62,41 / 814 /T) | synonymous_variant | LOW | 669C>T | 1497 | C223C | Only on D701N cluster sequences |
| OQ352551.1 / Seg 7 (M) | 691 | T | C | 47,79 / 15659 (C) | 59,61 / 3442 (C) | 59,31 /3728 (C) | synonymous_variant | LOW | 666T>C | 759 | H222H | Only on D701N cluster sequences |
|  | 691 | T | C | 47,79 / 15659 (C) | 59,61 / 3422 (C) | 59,31 /3728 (C) | intron_variant | MODIFIER | 27-49T>C | 294 |  | Only on D701N cluster sequences |
| OQ352552.1 / Seg 8 (NS) | 102 | G | A | 52,86 / 32303 (A) | 64,3 / 372 (A) | 62,81 / 5230 (A) | intron_variant | MODIFIER | 30+46G>A | 366 |  | Present on 44 sequences |
|  | 703 | T | C | 70,9 / 33304 (C) | 83 / 3700 (C) | 82,62 / 5312 (C) | synonymous_variant | LOW | 205T>C | 366 | L69L | Present on D701 cluster and one more Se lion sequence |
|  | 102 | G | A | 52,86 / 32302 (A) | 64,3 / 3712 (A) | 62,81 / 5230 (A) | missense_variant | MODERATE | 76G>A | 693 | E26K | Present on 44 sequences |
|  | 703 | T | C | 70,9 / 33304 (C) | 83 / 3700 (C) | 82,62 / 5312 (C) | missense_variant | MODERATE | 677T>C | 693 | I226T | Present on D701 cluster and Sea lion sequences |

## Discussion

The emergence and evolution of HPAIV H5N1 in Chile since December 2022 has significant implications for animal and public health. A collaborative effort between public and academic institutions between December 2022 and March 2023 procured over 4,000 samples for HPAIV H5N1 identification by active surveillance to monitor the outbreak and limit the spillover events to other hosts. The high number of samples from poultry, wild birds, and marine mammals provided valuable insights into the spread and impact of the H5N1 outbreak in Chile, informing targeted strategies for disease management and control in both wildlife and livestock populations. Genomic analysis provided insights into the rapid evolution and spatial dissemination of viruses with new genetic traits of concern. The identification of a positive human case early on during the outbreak underscores the zoonotic risk associated with HPAIV H5N1 and the need to continue monitoring the fast-changing evolutionary trajectory of the HPAIV outbreak in Chile.

The widespread detection of HPAIV in wild birds (18.5% positivity rate) indicates a high presence of the virus in avian populations. Most samples were collected from swabs taken from deceased animals in areas where HPAIV H5N1 had been detected. Some of them belonged to mass mortality events, especially for pelicans, shags, sea lions, and penguins. The multispecies detection of the HPAIV H5N1 2.3.4.4b lineage, including different orders of birds and mammals, highlights the potential for cross-species transmission and mammalian adaptation of the virus. Prior to the outbreak reported here, Chile’s poultry was considered free of avian influenza, and vaccination is not allowed due to commercial restrictions ^43^. Thus, surveillance and biosecurity measures have been reinforced to control the spread of HPAIV H5N1 in domestic birds and assess and reduce the risk of zoonotic transmission. The current outbreak has resulted in 160 positive backyard farms and 12 poultry farms that have been confirmed as HPAIV H5 positive, resulting in about 1,4 million poultry animals being culled or have died due to the current outbreak.

Phylogenetic analysis showed that all eight segments of the H5N1 genomes fall into a unique Peru/Chile clade, indicating a single introduction from North America. So far, no reassortment was observed in the cases analyzed here. This lack of reassortment could be explained by several factors such as the severity of the HPAIV H5N1 infection causing rapid death, a low prevalence of other AIV subtypes infecting the same avian population, or a host-related reduced viral-mixing capacity ^44^. Prior surveillance studies have detected endemic AIVs circulating in Chile’s wild aquatic birds that belong to the dominant lineages in the Americas and Eurasia, as well as a South America-specific lineage that has only been found in Chile, Argentina, and occasionally in Peru ^45–47^. While HPAIV H5N1 has reassorted frequently with the dominant American lineage in North American birds, producing multiple new genotypes, thus far there has been no reassortment between HPAIV and the South American lineage in Chile or any other sampled South American country. Current data indicates that multiple independent introductions of HPAIV H5N1 have occurred from North America into other parts of South America (Venezuela, Ecuador, Colombia), which are distinct from the introduction identified in Peru/Chile ^6, 12^. There are no signs of onward transmission to additional countries in South America, with the exemption of the limited H5N1 data from Argentina (one full-length sequence), which positioned this isolate in the Chile/Peru clade, suggesting the spread from Chile/Peru to Argentina possibly through the southern tip of the continent (Patagonia). Nonetheless, additional data from Argentina is needed to elucidate the predominant lineages circulating in wild animals.

Adaptation of the HPAIV H5N1 viruses to mammals is a significant pandemic concern due to its potential for zoonotic transmission, causing severe illness and the possibility of the virus to acquire human-to-human transmission. In the HPAIV H5N1 outbreak in Chile, we identified the D701N and Q591K PB2 mutations, in four animals and a human, which have been associated with mammalian adaptation. Data from current and previous HPAIV H5 outbreaks in Asia, Europe, and North America indicate that both mutations are rare ^48, 49^. Notably, we found in four viruses in Chile, including two sea lion viruses, one human case, and one sanderling carrying the D701N, and the Q591K mutation was observed in the human case and one of the sea lions. The D701N mutation has been shown to enhance viral replication and pathogenicity in mammalian hosts, including humans ^49–51^. The Q591K mutation has also been implicated in increased replication and transmission of the virus in mammals ^52, 53^. The D701N mutation was first reported in South America earlier in February (2023) in a Sea lion in Southern Peru. Our analysis suggests that there was no onward transmission of this particular virus to other hosts. Instead, a cluster of Chile/Peru viruses with the PB2 D701N mutation are from March 2023 and are positioned in a different section of the tree, along the human sequence. The D701N mutation was also reported on the Maine seal outbreak and on occasional infections of mammals in the USA, Canada, Finland, and Belgium. However, in these cases there was no clustering of mutant cases or signs of onward transmission.

In Chile, it remains less clear whether any transmission of D701N mutants occurred among the avian (sanderling) and/or mammalian (sea lion, human) hosts in the Chile/Peru D701N cluster. In **Figure 3B**, we outline three possible scenarios that could explain the apparent clustering of D701N (and Q591K) mutants in our dataset, while taking into account missing data. In scenario one (‘sea lion transmission’), viruses with D701N successfully transmit from sea lion to sea lion and occasionally spillover from sea lions to humans and birds. In scenario two (‘avian transmission’), viruses with D701N mutation transmit bird-to-bird and occasionally spillover from birds to humans and sea lions. In scenario three (‘no transmission, *de novo* mutation’), viruses with D701N are never transmitted between any hosts. Instead, multiple within-host *de novo* mutation events occur independently in human and animal hosts. In this case, the D701N viruses are not a “true” transmission cluster but rather cluster together due to a sampling artifact and the large amount of missing data from unsampled viruses from the Chilean outbreak.

Going forward, as more sequence data becomes available from the Chilean outbreak in the future, it is possible we would uncover more viruses positioned in this cluster that lack the D701N mutation, confirming scenario three (‘no transmission’). The presence of a within-host mixture of sequences with and without the D701N mutation also could support scenario three, if these mixed viral populations represent *de novo* within-host mutation events that occurred independently in each host. Alternatively, a loose transmission bottleneck could allow mixed viral populations to be transmitted between hosts. Continued surveillance and monitoring of the HPAIV outbreak in Chile, combined with experimental studies using animal models to monitor the phenotypic change, are needed to assess the risk posed by the H5N1 outbreak in Chile and inform public health measures to control its spread and protect human populations.

## Acknowledgments

We thank the Agricultural and Livestock Service (SAG) and SERNAPESCA personnel for their support and contributions, especially in the sample collection. We are grateful to Belen Aguero and Vanessa Mendieta from the Animal Virology Lab, Universidad de Chile, for their help in sample processing. Thanks to the Unidad de Bioinformática y Biología Computacional Pontificia Universidad Católica de Chile for his support on the bioinformatics pipeline. We are grateful to the GISAID EpiFlu™ Database, laboratories, and source of original data of influenza A virus (IAV) sequences, especially to the Servicio Nacional de Sanidad Agraria del Perú – SENASA, Peru, source of the strain closest strains and the Institute of Public Health of Chile. Thanks to Alonso Parra from MINSAL for valuable input about the human case. This work was also supported by the Centers of Excellence for Influenza Research and Response, National Institute of Allergy and Infectious Diseases, National Institutes of Health (NIH), Department of Health and Human Services, under contract 75N93021C00014, and by the Intramural Research Program of the US National Library of Medicine at the NIH. The funders had no role in study design, sample collection, data collection and analysis, decision to publish, or preparation of the article.

---

**Supplementary Table 1**. Summary of HPAIV animal cases according to host orders from December 09, 2022, to March 14, 2023, in Chile.

**Supplementary Table 2.** Summary of sequencing results of the surveillance of highly pathogenic avian influenza virus H5N1 in Chile. December 09, 2022, to March 14, 2023, in Chile.

**Supplementary Table 3.** Summary complete list of positive cases.

**Supplementary Table 4.** Summary of the minor variant analysis of the Chilean HPAIV genomes. Include SNV, single nucleotide variants; indels, insertions and deletions.

**Supplementary Table 5.** Complete list of the HPAIV variations classified according to the allele effect.

**Supplementary Figure 1:**
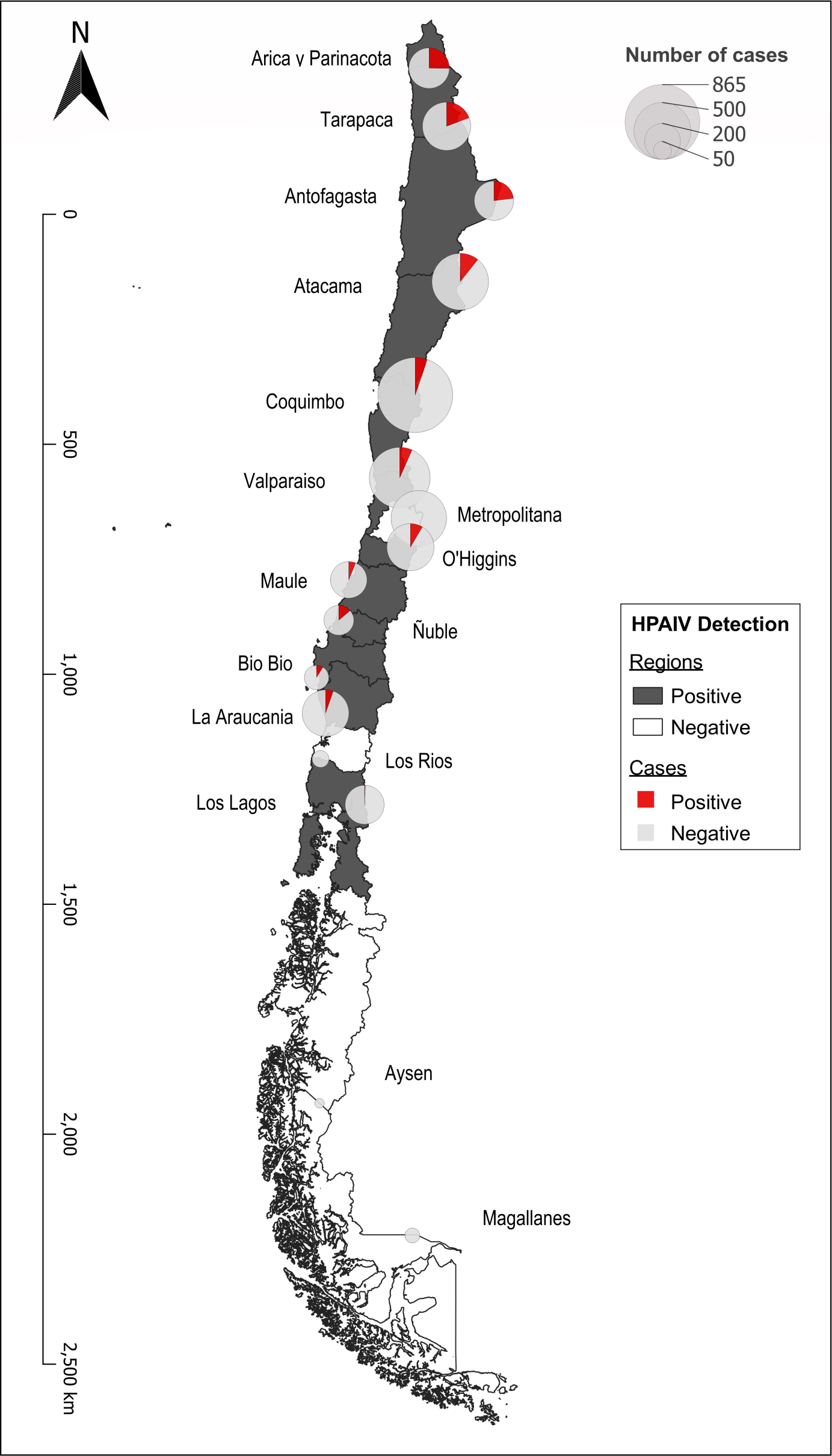
Map of Chile indicating HPAIV-positive cases by region between 12-toco09-2022 to 03-14-2023.

